# Subfunctionalization versus neofunctionalization after whole-genome duplication

**DOI:** 10.1101/190090

**Authors:** Simen R. Sandve, Rori V. Rohlfs, Torgeir R. Hvidsten

**Affiliations:** Center for Integrative Genetics (CIGENE), Faculty of Biosciences, Department of Animal and Aquacultural Sciences, Norwegian University of Life Sciences, Ås, Norway; Department of Biology, San Francisco State University, San Francisco, California, USA; Faculty of Chemistry, Biotechnology and Food Science, Norwegian University of Life Sciences, Ås, Norway; Umeå Plant Science Centre, Department of Plant Physiology, Umeå University, Umeå, Sweden

## Abstract

The question of what is the predominant evolutionary fate of genes after duplication events has been hotly debated for decades^1,2^. Two recently published articles in Nature (Lien *et al*.^3^) and Nature Genetics (Braasch *et al*.^4^) investigated the regulatory fate of gene duplicates after the salmonid-specific (Ss4R) and teleost specific (Ts3R) whole genome duplication (WGD) events, respectively. Both studies relied on tissue expression atlases for estimating regulatory divergence and used closely related unduplicated sister taxa (i.e. Northern pike and the spotted gar, respectively) as proxies for the ancestral expression state. Surprisingly, the two studies reach very different conclusions about the evolutionary mechanisms impacting gene expression after WGD. Braasch *et al*.^4^ concluded that the expression divergence was consistent with partitioning of tissue regulation between duplicates (subfunctionalization), while Lien *et al*.^3^ concluded that most divergence in tissue regulation were consistent with one copy maintaining ancestral tissue regulation while the other having diverged (in line with neofunctionalization). Here we show that this striking discrepancy in the conclusions of the two studies is a consequence of the data analysis approaches used, and is not related to underlying differences in the data.

---

To evaluate the underlying cause of the discrepancies between the two studies, we re-analysed the data from Braasch *et al*.^4^ using the approach of Lien *et al*.^3^, and vice versa. Both studies computed expression correlation to the unduplicated sister taxa as a measure of divergence - for the two duplicates individually (*Duplicate 1* and *Duplicate 2*) and for the summed expression of the duplicates (*Sum duplicates*). The only aspect of the analyses that differed was how the individual genes within the duplicate pairs were ranked. In Braasch *et al.*^4^, the duplicate pairs were ordered randomly (i.e. the two genes were assigned labels *Duplicate 1* and *Duplicate 2* at random, see Chapter 13.1, p. 36 of the supplementary materials of Braasch *et al.*^4^). In contrast, Lien *et al*.^3^ ranked the two genes in each duplicate pair as ‘most diverged’ (i.e. the gene with the lowest expression correlation to the ortholog in the unduplicated sister taxa, *Duplicate 1*) and ‘most conserved’ (i.e. the gene with the highest expression correlation to the ortholog, *Duplicate 2*).

**Figure 1:**
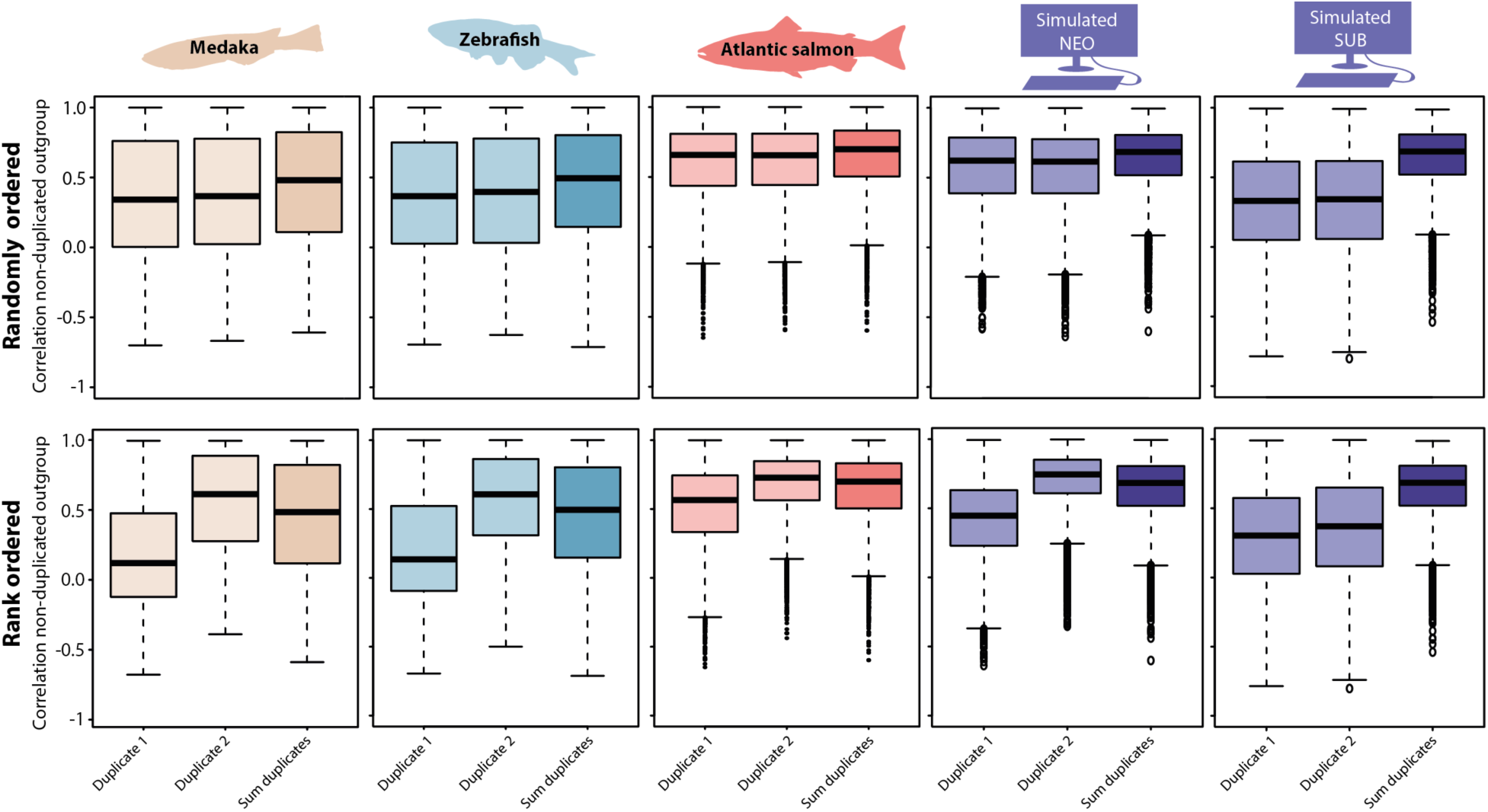
Tissue expression divergence in real and simulated data. Tissue expression correlation between duplicates in Medaka/Zebrafish and the corresponding orthologs in Spotted gar (1606 and 1315 triplets, respectively) and between duplicates in Atlantic salmon and the orthologs in Northern pike (8070 triplets). In the upper row, duplicated genes are assigned labels Duplicate 1 and Duplicate 2 randomly, while in the lower row the duplicates are ranked so that Duplicate 1 has the lowest correlation to the ortholog and Duplicate 2 the highest. Sum duplicates represents the correlations between the summed expression of the two duplicates and the ortholog in the unduplicated species. All correlations were computed using the Pearson correlation coefficient on the original expression data from the two publications in the first three columns and simulated data in the two last columns. All pairwise comparisons were statistically significant (p-value < 5e-11, Wilcoxon signed rank test, two-sided), with the exception of comparisons between Duplicate 1 and Duplicate 2 in the upper row (randomly ordered). The expression data is available in Supplementary Data Set 1.

Based on the three distributions of expression correlations between the ortholog in the unduplicated sister taxa (i.e. assumed ancestral state) and *Duplicate 1*, *Duplicate 2* and *Sum duplicates* in the duplicated species, we can draw conclusions about broad transcriptome-wide evolutionary trends. If subfunctionalization is the dominating driver of expression divergence, we expect the sum-of-duplicates to correlate more strongly with the ancestral state than the individual gene duplicates. Conversely, if neo-functionalization is the major evolutionary mechanism in play, we expect that the duplicate with the most conserved expression patterns (*Duplicate 2* in the method that rank duplicates) to exhibit the highest expression correlation.

Our re-analyses of both datasets show that the conclusions from the two papers would be identical if the same data analysis approach were used (Figure 1). So where does this leave us? Do the Braasch *et al*.^4^ and Lien *et* al.^3^ results support sub-or neofunctionalization as the dominating mechanism?

To answer this question, we simulated neo-and subfunctionalized duplicates and applied both the randomly ordered and rank ordered methods. For the simulations, we use the pike transcriptome as an unduplicated reference. For the duplicates, we simulate neofunctionalization by adding a small error term ∼N(0,1) for the conserved duplicate, and a larger error term ∼N(0,5) for the diverged duplicate. Sub-functionalization was simulated by randomly partitioning the pre-duplicated tissue expression across the two duplicates, and then adding a small error term ∼N(0,1). While the ranked approach correctly identified patterns consistent with sub-and neofunctionalization in the simulated data (Figure 1, bottom row), random ordering of duplicates obscures the signal of neofunctionalization, and exhibited patterns consistent with subfunctionalization for both simulations (Figure 1, top row).

From this evidence, we conclude that the prevalent fate of gene duplicates from Ts3R and Ss4R WGD is that one duplicate is under stronger purifying selection pressure to maintain ancestral regulation than the other duplicate. This is more consistent with regulatory neofunctionalization than with subfunctionalization. Nevertheless, the observed asymmetrical gene expression divergence among duplicates could be a result of relaxed purifying selection (i.e. neutral evolution) rather than adaptive selection for a new regulation (i.e. neofunctionalization). This crux can be addressed in future studies by comparing the likelihood of the observed duplicate expression divergence data across multiple species under a model of regulatory neofunctionalization and under a model of neutral evolution (see for example ref. 8). Such a test will have improved power when including more informative species and will provide strongest evidence for neofunctionalization when duplicate divergence is conserved across several species sharing the WGD. Finally, by only evaluating global expression evolution patterns we neglect that selection act differently on different individual genes. It is therefore of the utmost importance to adapt existing phylogenetic methods^5-8^ and to develop new techniques to evaluate whether the data supports sub-or neofunctionalization on a gene-by-gene basis. Now that genome sequences and associated functional data is increasingly accumulating, this will open up new and exciting avenues for answering long standing questions regarding genome evolution following whole genome duplication.

## Methods

Expression data for gene triplets (i.e. *Duplicate 1* and *Duplicate 2* in the duplicated species and their ortholog in the unduplicated sister species) were taken from Lien *et al*.^3^ and Braasch *et al*.^4^ (Supplementary Data Set 1). This included three RNA sequencing data set: Medaka/Spotted gar (1606 triplets, 11 tissues), Medaka/Spotted gar (1606 triplets, 11 tissues) and Atlantic salmon/Northern pike (8070 triplets, 13 tissues). We also generated data by simulating the processes of neo-and subfunctionalization. In these simulations, the Northern pike data set was used as the unduplicated reference. To simulate the process of neofunctionalization, we duplicated each gene in the reference data set and added a small error term ∼N(0,1) to one (conserved duplicate) and a larger error term ∼N(0,5) to the other (diverged duplicate). Subfunctionalization was simulated by randomly partitioning the tissue expression values of each gene in the unduplicated reference across two duplicates. Specifically, for each reference gene we (i) randomly sampled the tissues (out of 13) that would be expressed in each duplicate, and (ii) assigned the corresponding reference expression values to the simulated duplicates. The tissues in the duplicates that were not assigned expression values were set to zero, hence the expression of the duplicates summed to that of the unduplicated reference. Finally, a small error term ∼N(0,1) was added to all expression levels.

Pearson correlations were computed between the expression profile of each duplicate (*Duplicate 1* and *Duplicate 2*) and the ortholog as well as between the summed expression of the duplicates (*Sum duplicates*) and the ortholog. The correlations from each data set was then visualised using boxplots (*boxplot*-function in R using default settings), with boxes for *Duplicate 1*, *Duplicate 2* and *Sum duplicates*. Two different methods for identifying genome wide signatures of neo-and subfunctionalization was applied. One were the duplicates were named *Duplicate 1* and *Duplicate 2* by random (Randomly ordered, Braasch *et al*.^4^) and one were the duplicate with the lowest correlation to the ortholog in the unduplicated species was named *Duplicate 1* and the duplicate with the highest correlation to the ortholog was named *Duplicate 2* (Rank ordered, Lien *et al*.^3^). All pairwise comparisons of correlations within one dataset were statistically significant (p-value < 5e-11, Wilcoxon signed rank test, two-sided, *wilcox.test-*function in R), with the exception of comparisons between *Duplicate 1* and *Duplicate 2* when these were ordered randomly.

## Acknowledgements

S.R.S was supported by the Norwegian Research Council projects 221734 and 244164.

## Author contributions

S.R.S. and T.R.H. conceived the study. T.R.H. analysed the expression data, and R.V.R. performed the simulations. All authors interpreted the results, and wrote and approved the manuscript for publication.

